# Development and Validation of a Comprehensive Tumor Treating Fields System for Glioblastoma Therapy: From Prototype Design to Preclinical Evaluation

**DOI:** 10.1101/2024.10.14.618144

**Authors:** Xindong Wang, Han Lv, Zhiyong Wang, Xian Wang

**Affiliations:** Institute of Flexible Electronics Technology of THU, Zhejiang, Jiaxing, 314006, China; Sun Yat-Sen Memorial Hospital, Sun Yat-Sen University, Guangzhou, 510120, China; Guangzhou Medical University, Guangzhou, 511436, China

**Keywords:** Glioblastoma, Tumor Treating Fields, Electric Field Therapy, System Design, TTF System

## Abstract

Glioblastoma (GBM) is an aggressive brain tumor with limited treatment options and a poor prognosis. Tumor Treating Fields (TTF) therapy, which uses alternating electric fields to disrupt cancer cell division, has emerged as a promising non-invasive treatment. However, the development and optimization of TTF systems remain challenging, with limited studies detailing the design and practical application of such systems. In this study, we developed a novel TTF prototype, focusing on the design and fabrication of key components such as the electrical signal regulation, transmission and corresponding therapeutic effects. We evaluated the system’s efficacy through a series of *in vitro* and *in vivo* experiments. *In vitro*, we demonstrated significant inhibition of GBM cell proliferation under varying electric field intensities, with stronger fields showing greater efficacy. *In vivo* studies using a rat glioblastoma model revealed reduced tumor growth, increased apoptosis, and enhanced immune infiltration in the TTF-treated group compared to controls. This comprehensive study provides a valuable reference for TTF system development, offering insights into both the technical design and biological application of TTF therapy, and highlighting its potential for improving GBM treatment outcomes.

## Introduction

Glioblastoma (GBM) is the most aggressive and common form of primary brain tumor in adults, representing approximately 15% of all brain tumors[1, 2]. Despite aggressive treatment approaches including surgery, radiation, and chemotherapy, the prognosis for GBM patients remains dismal, with a median survival time of only 12 to 15 months[3]. The challenges in treating GBM stem from its highly invasive nature, rapid proliferation, and resistance to standard therapies. Current therapies are often limited by the blood-brain barrier, which restricts the delivery of chemotherapeutic agents, as well as the tumor’s heterogeneity, which leads to treatment failure and recurrence[4]. Given the poor outcomes associated with existing treatment regimens, there is a critical need for novel therapeutic strategies that can target GBM more effectively[5, 6].

Tumor Treating Fields (TTF) have emerged as a promising non-invasive therapeutic modality for GBM, offering an alternative mechanism to target tumor growth[7, 8]. TTF utilizes low-intensity, alternating electric fields at intermediate frequencies (100-300 kHz) to disrupt the mitotic process of rapidly dividing tumor cells. These fields interfere with microtubule polymerization, alignment of chromosomes, and other essential mitotic functions, leading to apoptosis[9, 10]. The U.S. Food and Drug Administration (FDA) has approved the use of TTF therapy through devices like the Optune system, further validating its potential in GBM treatment. Clinical trials have demonstrated the ability of TTF therapy to prolong overall survival and progression-free survival in GBM patients when combined with standard chemoradiation[11, 12]. However, the full potential of TTF has yet to be realized, as current systems are limited in their flexibility to adjust field parameters[11, 13, 14], and few studies have explored the detailed engineering behind the design and optimization of TTF systems.

To address these limitations, our study presents the design, development, and preclinical evaluation of a novel TTF system, focusing on key components such as the electrical signal regulation, transmission and corresponding therapeutic effects. We designed the system to generate highly controlled electric fields and incorporated high-dielectric ceramic electrodes composed of barium titanate zirconate, which have superior electric field transmission properties compared to conventional electrodes. By evaluating the performance of this system through *in vitro* glioblastoma cell experiments and *in vivo* rat models, we demonstrate its efficacy in inhibiting tumor growth and inducing apoptosis. Furthermore, we provide a comprehensive guide to the technical design of the system, aiming to make TTF research more accessible and encourage further innovation in the field. This study contributes valuable insights into the optimization of TTF therapy for clinical use, particularly in enhancing the effectiveness of electric field-based cancer treatments.

## Materials and Methods

### 1. Reagents and Instruments

The reagents used in the experiments comprised Isoflurane (RWD Life Science Co., Ltd.), D-Luciferin Potassium Salt (Rhone Reagents), glioblastoma cells and C6-LUC glioma cells (Cyagen Biosciences), DMEM medium (Procell Life Science & Technology Co., Ltd.), Penicillin-Streptomycin (Procell Life Science & Technology Co., Ltd.), and Fetal Bovine Serum (ExCell Bio). Staining reagents included Hematoxylin Stain Solution (Zhuhai Baso Biotechnology Co., Ltd.), Eosin Y (Sinopharm Chemical Reagent Co., Ltd.), Ethanol (Sinopharm Chemical Reagent Co., Ltd.), Dewaxing Agent (Zhuhai Baso Biotechnology Co., Ltd.), Paraffin (Leica Biosystems), Hydrochloric Acid, Ammonia Solution (Sinopharm Chemical Reagent Co., Ltd.), and Mounting Medium (Zhuhai Baso Biotechnology Co., Ltd.). Antibodies used for immunohistochemistry included Ki-67 (Abcam), CD8 (Boster Bio), and Caspase-3 (Abcam). The ceramic electrodes were from the Flexible Bioelectronics Division, Institute of Flexible Electronics Technology of THU. Sprague-Dawley (SD) rats were used in the experiments, sourced from Hangzhou Qizhen Experimental Animal Technology Co., Ltd. All rats were male, aged between 6-8 weeks.

The primary instruments used in this study included a Bruker D8 Discover X-ray diffractometer (XRD), an Agilent 4294A precision impedance analyzer, a Zeiss Sigma 300 scanning electron microscope (SEM), a stereotaxic apparatus (Blue Star B/S, Anhui Zhenghua Biological Instrument Co., Ltd.), a multimode small animal *in vivo* imaging system (AniView 600, Guangzhou Boluoteng Biotechnology Co., Ltd.), a small animal anesthesia machine (Model H1670401-200L, RWD Life Science Co., Ltd.), and a small animal magnetic resonance imaging (MRI) system (7.0T MRI, Bruker Biospin GmbH PharmaScan7016). Additional equipment included a dehydrator (Excelsior AS, Thermo Fisher Scientific), an embedding machine (Model YB-6LF, Xiaogan Yaguang), a microtome (Model HM 340E, Thermo Fisher Scientific), a constant temperature drying oven (Model DHP-9082, Ningguo Shaying Scientific Instrument Co., Ltd.), a slide scanner (Pannoramic MIDI, 3D), and an automatic staining machine (Gemini AS, Thermo Fisher Scientific). A dehydrator and embedding machine were used for histopathological sample processing, and a slide scanner was employed for imaging the stained samples.

### 2. Cell Experiments

Four round glass slides were placed in the TTF cell experimental device (Figure 1c and 2a). 3 mL of cell suspension (density 2×10^5^ cells/mL) was applied evenly on a slide, and incubated at 37 °C. The TTF-treated group was subjected to the alternating electric field with specific parameters (field strength and frequency). Control cells received no electric field stimulation but were otherwise cultured under identical conditions. The slides were taken out at specified times, and the cells were digested and counted. Cell morphology and proliferation rates were recorded at each time point, and changes in cell density were analyzed to assess the effect of TTF on cell growth.

**Figure 1.**
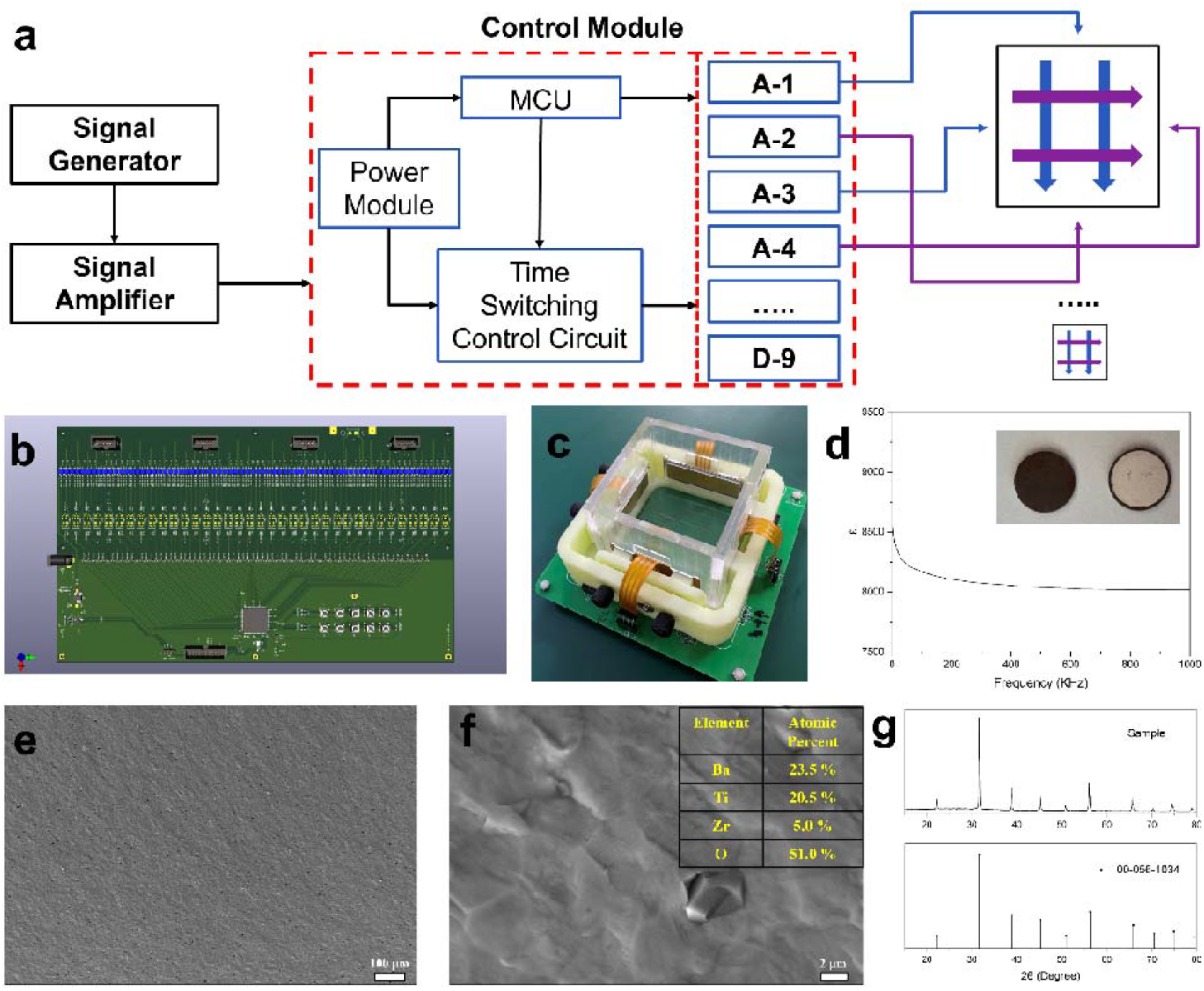
(a) The overall design block diagram of the TTF system. (b) The PCB layout of the Control Module. (c) An actual photograph of the TTF system used in the cell experiments. (d) The dielectric constant measurements of the ceramic electrode across different frequencies, the inset shows the front and back of a ceramic electrode, with the back silver plated. (e, f) SEM images and EDS analysis showing the cross-sectional morphology of the ceramic electrode. (g) The XRD pattern of the ceramic electrode.

### 3. Rat Tumor Model

Rats were acclimatized in an SPF-grade environment at a controlled temperature of 20-26°C and humidity of 40-70%, with a 12-hour light-dark cycle. The rats were provided sterilized corn cob bedding, along with free access to food and water for 7 days. Following acclimation, the rats were divided into two groups: the control group and the TTF treatment group, with three rats in each group.

The glioblastoma model[15, 16] was established by stereotaxic injection of C6-LUC cells. Rats were anesthetized using isoflurane and secured in a stereotaxic frame. The C6-LUC cells, harvested during their exponential growth phase, were prepared at a concentration of 5×10^5^ cells in 5 µL PBS. The cells were injected into the right striatum at coordinates 1 mm posterior to bregma, 3 mm lateral to the midline, and 4.5 mm depth from the skull surface. After injection, the skull was sealed with bone wax, and the incision was sutured and disinfected. The rats were then returned to the SPF environment for recovery.

### 4. TTF Treatment

Three days post-injection, TTF treatment commenced. Anesthetized rats had electrode patches affixed to their shaved scalp, positioned orthogonally to create intersecting electric fields. The TTF system was activated for >20 hours daily, with continuous treatment until day 20. The electric field parameters were set to 2 V/cm at 200 kHz, matching the in vitro settings. Every four days, the electrodes were removed, the skin was disinfected, and fresh electrodes were attached.

Rats were monitored daily for general health, and the skin reactions at the electrode sites were recorded. Bioluminescent imaging was conducted on days 10, 12, 14, 16, and 18 using an *in vivo* imaging system following intraperitoneal injection of D-luciferin. MRI scans were performed on day 19 using a T2-TurboRARE sequence to confirm tumor size. On day 19, blood samples were collected for complete blood count and biochemical analysis. Brain tissues were fixed, embedded in paraffin, and sectioned for immunohistochemical analysis with antibodies against Ki-67, CD8, and Caspase-3.

Brain tissues were stained with hematoxylin and eosin (H&E) following standard protocols. For immunohistochemistry, sections underwent antigen retrieval using a citrate buffer, followed by incubation with primary antibodies at 4°C overnight. After incubation with secondary antibodies, DAB staining was used for visualization, and sections were counterstained with hematoxylin. Data from these histological analyses were used to assess tumor proliferation, immune infiltration, and apoptosis in response to TTF treatment.

## Results

The TTF system consists of a Signal Generator, Signal Amplifier, and Control Module (Figure 1a). The Signal Generator produces a 200 kHz sine wave, amplified by the Signal Amplifier, and modulated through a Microcontroller Unit (MCU)-controlled Control Module to adjust signal intensity and frequency for each of the 36 output channels. This modular design allows independent control over each electrode’s output, enabling precise application of electric fields. In the cell culture experiments, four electrodes are arranged orthogonally, providing alternating electric fields every second, as depicted in the inset of Figure 1a.

The control module’s PCB layout, shown in Figure 1b, highlights the intricate design needed to manage the precise delivery of the electric fields to each electrode. Figure 1c presents the prototype used in the cell experiments, demonstrating the physical implementation of the system with orthogonally positioned ceramic electrodes. A key feature of the system is the high-dielectric ceramic electrodes, which have a dielectric constant exceeding 8000 at 200 kHz, significantly higher than the reported electrodes[7] (Figure 1d). The superior dielectric properties ensure effective electric potential transmission during treatment. The electrode’s cross-sectional morphology, analyzed by the scanning electron microscope (SEM), reveals a uniform structure, and the energy dispersive spectrometer (EDS) confirms the primary composition as barium titanate zirconate (Figures 1e and 1f). The X-ray diffractometer (XRD) analysis (Figure 1g) further validates the crystalline structure of the ceramic electrodes, matching the standard PDF card 00-056-1034, confirming the suitability for stable and efficient electric field generation. The combination of these images confirms the successful design and fabrication of the TTF system, ensuring effective electric field delivery during therapy.

In the cell culture experiments, glioblastoma cells (U251) were exposed to varying electric field intensities using the TTF system. The experimental setup is illustrated in Figure 2a, where four circular cell culture slides were placed under orthogonally arranged electrodes to apply the electric fields in alternating directions. In Figure 2b, the electric field distribution across the culture area was simulated using the finite element analysis software (COMSOL Multiphysics), confirming the uniform application of the field over the cells.

**Figure 2.**
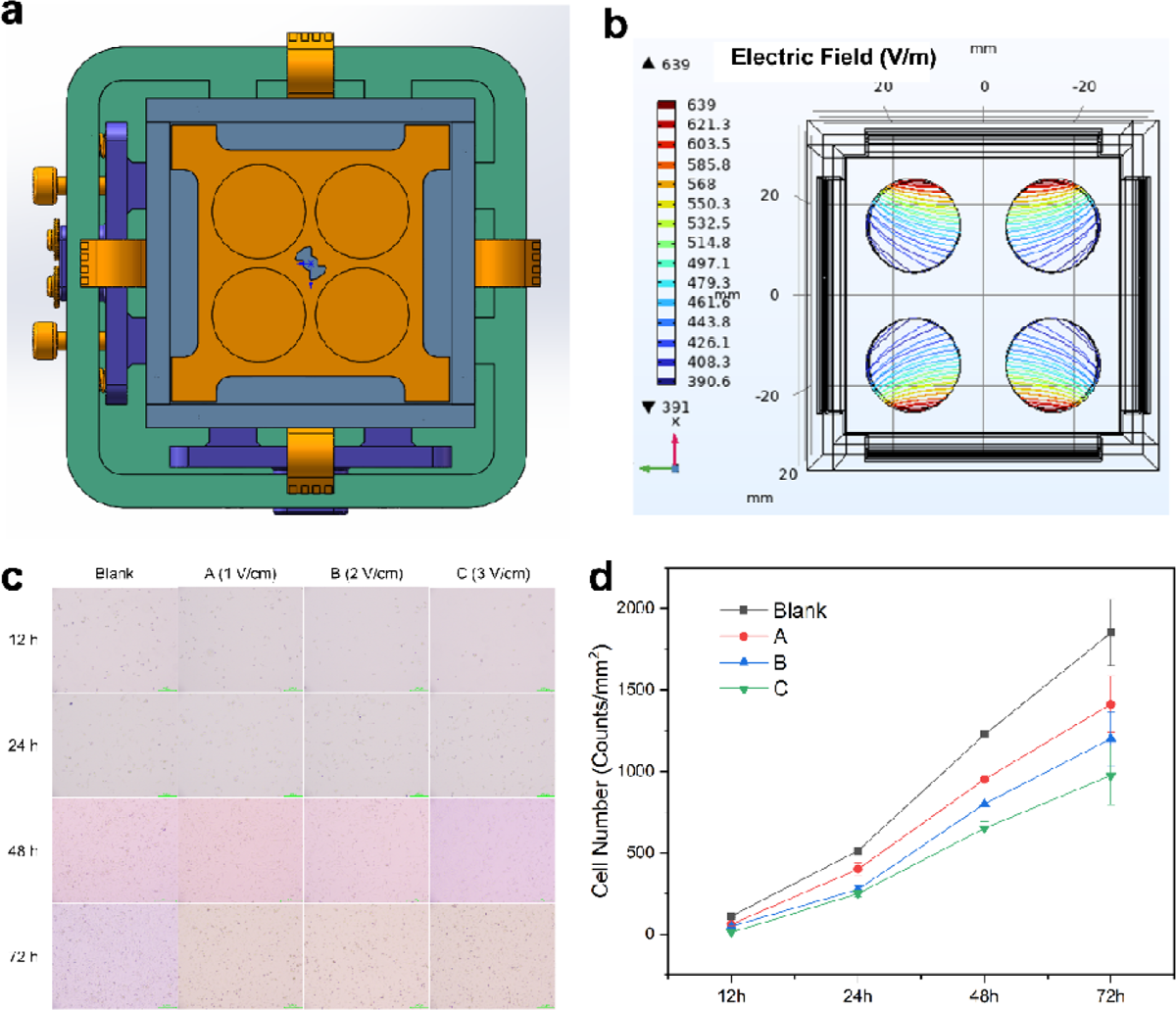
(a) The top view of the cell experiment prototype design, showing the placement of round cell culture glass slides and the orthogonal electric field application. (b) The finite element simulation of the electric field distribution using Comsol software. (c) Images of U251 cells cultured under different electric field intensities and at various time points (12, 24, 48, and 72 hours). (d) The quantitative analysis of adherent cell counts.

The visual comparison of cell morphology across different electric field strengths shows a significant reduction in cell proliferation, especially at higher field intensities. Figure 2c presents micrographs of cells treated under 1, 2, and 3 V/cm electric fields at 12, 24, 48, and 72-hour intervals. A marked decrease in cell number and altered cell morphology were observed at all time points in the TTF-treated group, with the highest intensity (3 V/cm) showing the most pronounced effects. After 72 hours, cell counts revealed a 35% reduction in proliferation at 2 V/cm and a 50% reduction at 3 V/cm compared to the control group (Figure 2d). These results demonstrate that the TTF system significantly inhibits glioblastoma cell growth *in vitro*. The inhibitory effect appears to be both time- and intensity-dependent, with higher electric field intensities resulting in greater inhibition of cell proliferation.

For the *in vivo* experiments, a rat glioblastoma model was established to evaluate the efficacy of TTF therapy. The ethical approval from the Ethics Review Committee was obtained prior to the research. Figure 3a shows the placement of four electrode patches on the rat’s head, positioned orthogonally to ensure comprehensive electric field coverage. C6-LUC glioblastoma cells, derived from the rat C6 glioma cell line by stably integration of a constitutive firefly luciferase stably expression construct, were injected into the brain, and TTF therapy was applied at 2 V/cm for >20 hours per day for 20 days.

**Figure 3.**
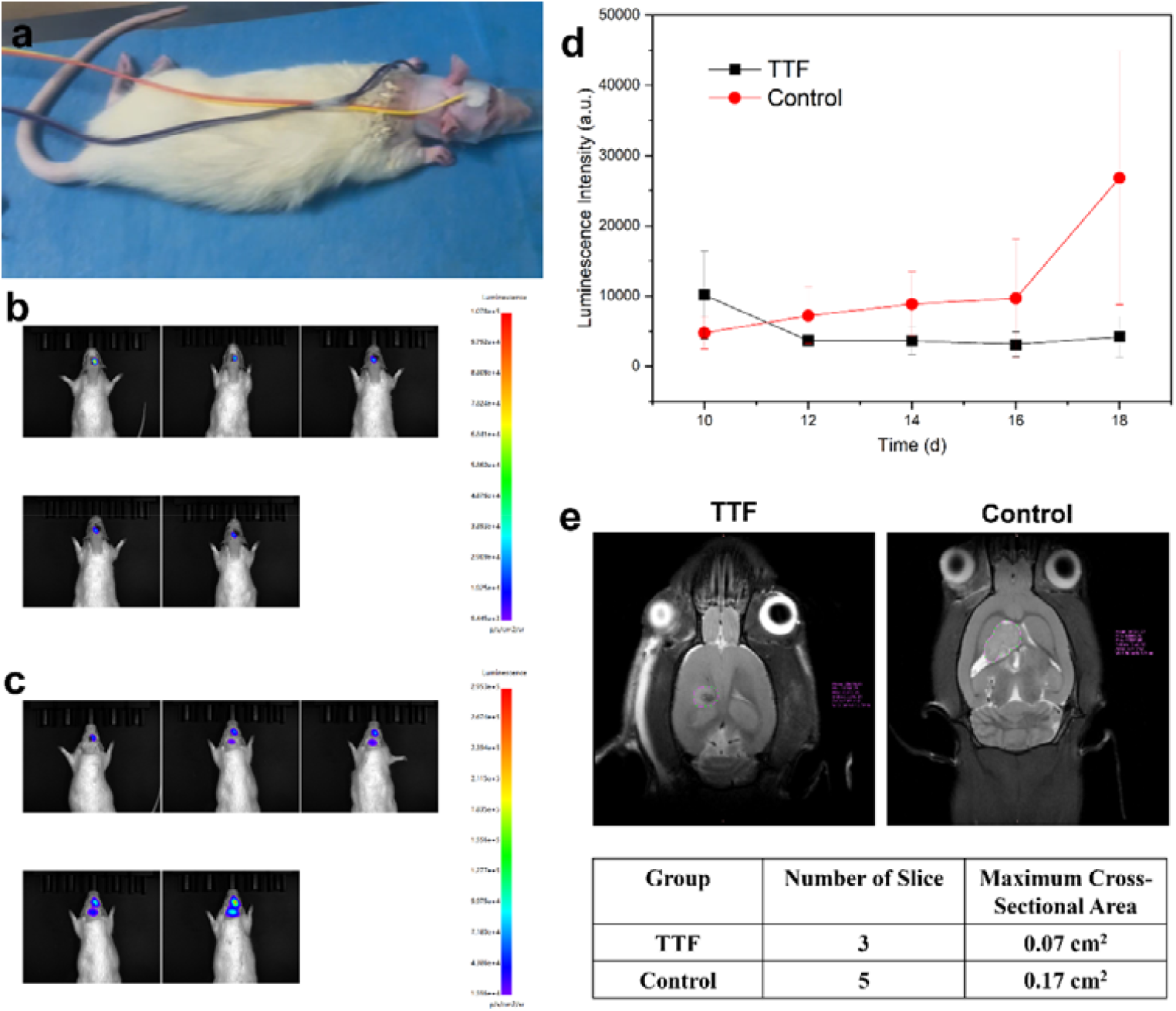
(a) The setup for the TTF treatment on the rat’s head with 4 electrode patches placed on both sides of the head. (b, c) The bioluminescence images of rats from the TTF-treated group and the control group, respectively, over several days. (d) The comparative graph of bioluminescence intensity between the treatment and control groups. (e) MRI scans from day 18 showing reduced tumor size in TTF-treated rats compared to the control group.

Tumor growth was monitored using bioluminescence imaging at multiple time points-days 10, 12, 14, 16, and 18 post-inoculation (Figures 3b and 3c). Bioluminescence intensity served as an indicator of tumor size, with stronger signals representing larger tumors. Comparison between the TTF-treated and control groups revealed significant tumor suppression in the treated group over time (Figure 3d). By day 18, tumor luminescence intensity was reduced by more than 5 times in the TTF-treated group compared to controls. Additionally, MRI scans performed on day 18 (Figure 3e) provided further validation of the tumor size reduction, with TTF-treated rats exhibiting smaller tumor cross-sectional areas compared to controls. These results demonstrate the effectiveness of the TTF system in slowing tumor progression in the glioblastoma rat model.

During TTF therapy, additional physiological parameters were monitored to assess treatment safety and tolerability in the glioblastoma rat model. Figure 4a shows the detailed process of electrode placement on the rats, ensuring proper alignment for optimal electric field delivery. Body weight changes were tracked throughout the experiment, as shown in Figure 4b, indicating that TTF-treated rats maintained a relatively stable body weight compared to the control group, suggesting minimal adverse effects on overall health. To assess potential thermal damage from the electrode patches, temperature measurements were recorded at the electrode-skin interface during TTF treatment (Figure 4c). The data showed no significant increase in skin temperature over time, confirming that the treatment did not induce thermal injury. However, some mild skin damage was observed at the electrode attachment sites on days 8, 10, and 12 (Figure 4d), where local redness and slight erosion occurred. These skin lesions were manageable and did not progress to severe injury, demonstrating that while there are some localized effects, the overall safety profile of the TTF system remains favorable.

**Figure 4.**
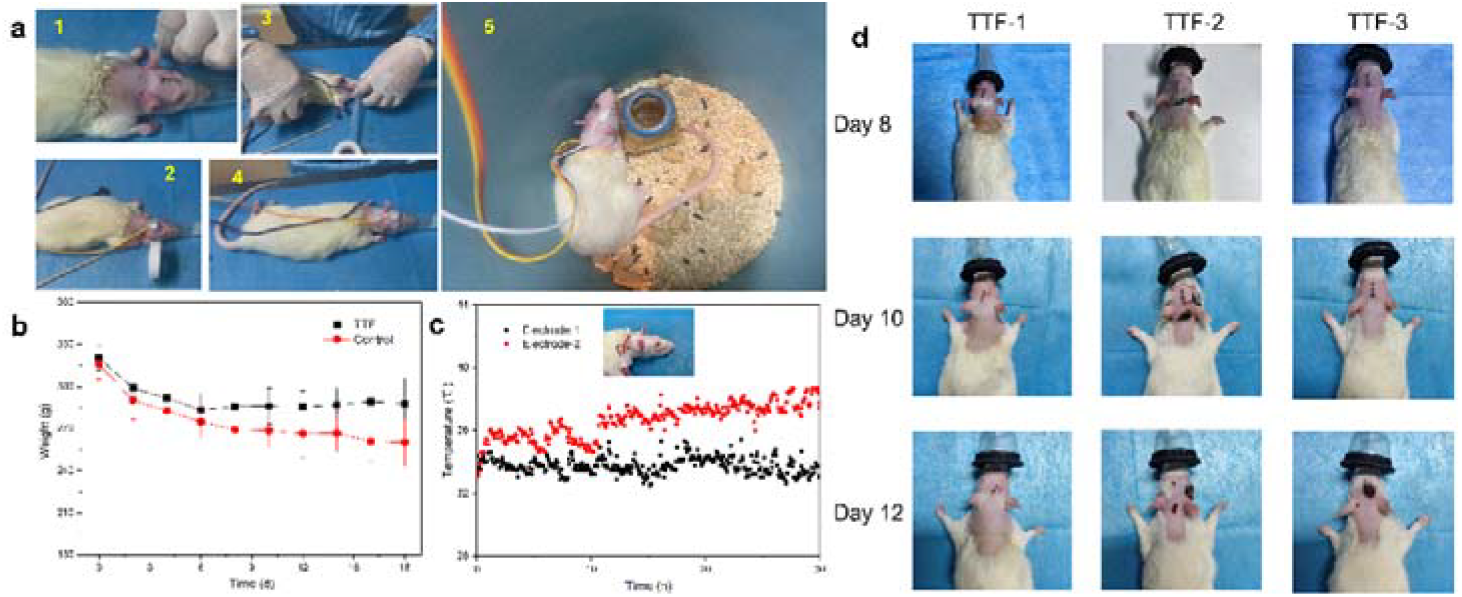
(a) The electrode installation process and treatment setup for rats undergoing TTF therapy. (b) The change in body weight over time for both control and TTF-treated groups. (c) The temperature profile at the electrode-skin interface during TTF therapy. (d) The skin condition of rats in the treatment group on days 8, 10, and 12.

Histopathological and immunohistochemical analyses were performed to assess the biological impact of TTF therapy on glioblastoma tumors[15, 16]. Figure 5a presents the H&E staining, as well as immunostaining for CD8, Caspase-3, and Ki-67, comparing the TTF-treated and control groups. In the H&E-stained sections, the TTF treatment group exhibited small, isolated tumor foci with sparsely arranged tumor cells and occasional mitotic figures, indicating a reduction in tumor cell density. In contrast, the control group showed large, dense tumor areas with abundant tumor cells, tightly packed arrangements, numerous blood vessels, and a high frequency of mitotic figures, highlighting the aggressive nature of the untreated tumors.

**Figure 5.**
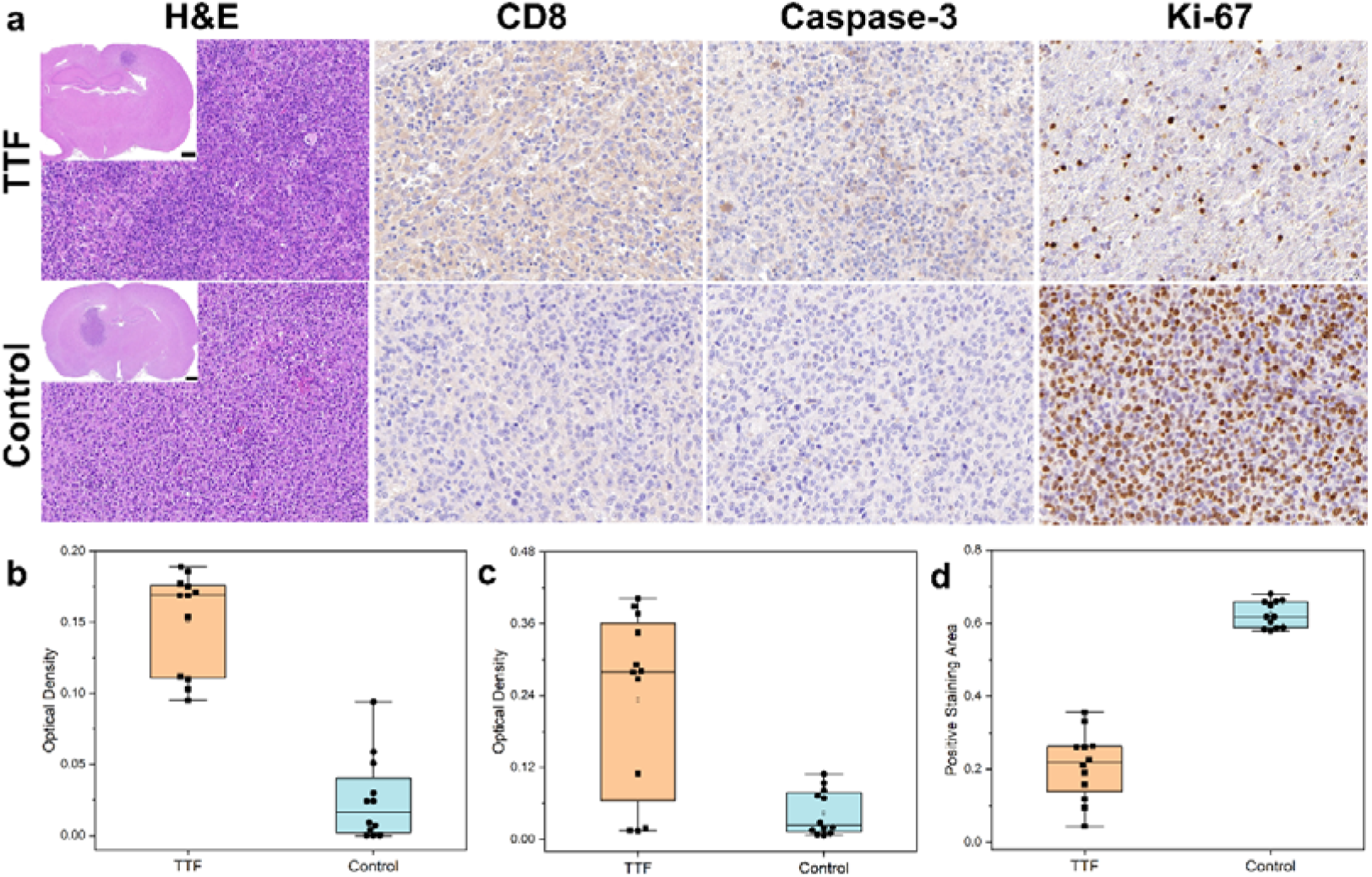
(a) The H&E, CD8, Caspase-3, and Ki-67 staining for both TTF-treated and control groups. (b) The quantitative analysis of immunoreactive cells, showing a significant increase in CD8 and Caspase-3 expression and a decrease in Ki-67 expression in the TTF group compared to controls.

The immunohistochemical staining further supports these findings. CD8 staining in the TTF-treated group showed increased infiltration of immune cells compared to the control group, suggesting an enhanced immune response (Figure 5a). Caspase-3 staining revealed elevated levels of apoptosis in the treatment group, confirming that the TTF therapy promoted tumor cell death. Conversely, Ki-67 staining, which marks proliferating cells, was significantly reduced in the TTF-treated group, indicating suppressed tumor cell proliferation. The quantitative data for CD8, Caspase-3, and Ki-67 positive cells are presented in Figures 5b, 5c, and 5d, respectively, demonstrating statistically significant differences between the treatment and control groups. These results confirm that TTF therapy not only inhibits tumor growth but also enhances apoptosis and stimulates immune activity.

## Conclusion

Our study presents a comprehensive and systematic approach to developing and evaluating a novel TTF therapy system, which successfully inhibits glioblastoma growth in both *in vitro* and *in vivo* models. By optimizing key system components, we demonstrated that TTF therapy effectively reduces tumor cell proliferation, enhances apoptosis, and may stimulate an anti-tumor immune response. Importantly, the system maintained an acceptable safety profile, with minimal adverse effects observed during treatment. These findings support the potential for TTF therapy to be further developed and optimized for clinical applications in glioblastoma and possibly other aggressive cancers.

Future directions for this research include exploring the potential combination of TTF therapy with chemotherapeutic agents or immunotherapies to enhance therapeutic efficacy. Additionally, further refinement of electric field parameters, such as frequency and duration, may help improve the effectiveness of this therapy in different cancer models.

## References

1. Yalamarty, S.S.K., et al., Mechanisms of Resistance and Current Treatment Options for Glioblastoma Multiforme (GBM). 2023. 15(7): p. 2116.

2. Saqib, M., et al., Clinical and translational advances in primary brain tumor therapy with a focus on glioblastoma-A comprehensive review of the literature. World Neurosurgery: X, 2024. 24: p. 100399.

3. Ostrom, Q.T., et al., CBTRUS Statistical Report: Primary brain and other central nervous system tumors diagnosed in the United States in 2010–2014. Neuro-Oncology, 2017. 19(suppl_5): p. v1–v88.

4. Wu, W., et al., Glioblastoma multiforme (GBM): An overview of current therapies and mechanisms of resistance. Pharmacological Research, 2021. 171: p. 105780.

5. Stupp, R., et al., Radiotherapy plus Concomitant and Adjuvant Temozolomide for Glioblastoma. 2005. 352(10): p. 987–996.

6. Davis, M.E., Glioblastoma: Overview of Disease and Treatment. Clin J Oncol Nurs, 2016. 20(5 Suppl): p. S2–8.

7. Kirson, E.D., et al., Alternating electric fields arrest cell proliferation in animal tumor models and human brain tumors. 2007. 104(24): p. 10152–10157.

8. Davies, A.M., U. Weinberg, and Y. Palti, Tumor treating fields: a new frontier in cancer therapy. 2013. 1291(1): p. 86–95.

9. Tuszynski, J.A., et al., An Overview of Sub-Cellular Mechanisms Involved in the Action of TTFields. 2016. 13(11): p. 1128.

10. Cao, Q., et al., Tumor Treating Fields (TTFields) combined with the drug repurposing approach CUSP9v3 induce metabolic reprogramming and synergistic anti-glioblastoma activity in vitro. British Journal of Cancer, 2024. 130(8): p. 1365–1376.

11. Stupp, R., et al., Maintenance Therapy With Tumor-Treating Fields Plus Temozolomide vs Temozolomide Alone for Glioblastoma: A Randomized Clinical Trial. JAMA, 2015. 314(23): p. 2535–2543.

12. Jackson, T.L., et al., A Randomized Controlled Trial of OPT-302, a VEGF-C/D Inhibitor for Neovascular Age-Related Macular Degeneration. Ophthalmology, 2023. 130(6): p. 588–597.

13. Rominiyi, O., et al., Tumour treating fields therapy for glioblastoma: current advances and future directions. British Journal of Cancer, 2021. 124(4): p. 697–709.

14. Guo, X., et al., Tumor-Treating Fields in Glioblastomas: Past, Present, and Future. 2022. 14(15): p. 3669.

15. Wu, H., et al., Exploring the efficacy of tumor electric field therapy against glioblastoma: An in vivo and in vitro study. 2021. 27(12): p. 1587–1604.

16. Wu, H., et al., Evaluation of a tumor electric field treatment system in a rat model of glioma. 2020. 26(11): p. 1168–1177.

